# Imprecise perception of hand position during early motor adaptation

**DOI:** 10.1101/2023.11.30.569377

**Authors:** Matthias Will, Max-Philipp Stenner

## Abstract

Localizing one’s body parts is important for movement control and motor learning. Recent studies have shown that the precision with which people localize their hand places constraints on motor adaptation. While these studies have assumed that hand localization remains equally precise across learning, we show that precision decreases rapidly during early motor learning. In three experiments, healthy young participants (n=92) repeatedly adapted to a 45° visuomotor rotation for a cycle of two to four reaches, followed by a cycle of two to four reaches with veridical feedback. Participants either used an aiming strategy that fully compensated for the rotation (experiment 1), or always aimed directly at the target, so that adaptation was implicit (experiment 2). We omitted visual feedback for the last reach of each cycle, after which participants localized their unseen hand. We observed an increase in the variability of angular localization errors when subjects used a strategy to counter the visuomotor rotation (experiment 1). This decrease in precision was less pronounced in the absence of re-aiming (experiment 2), and when subjects knew that they would have to localize their hand on the upcoming trial, and could thus fully allocate attention to hand position (experiment 3). We propose that attention to vision during strategic re-aiming decreases the precision of perceived hand position. We discuss how these dynamics in precision during early motor learning could impact on motor control, and shape the interplay between implicit and strategy-based motor adaptation.

**NEW & NOTEWORTHY:** Recent studies indicate that the precision with which people localize their hand limits implicit visuomotor learning. We found that localization precision is not static, but decreases early during learning. This decrease is pronounced when people apply a re-aiming strategy to compensate for a visuomotor perturbation, and diminishes when the hand is fully attended. We propose that these attention-dependent dynamics in position sense during learning may influence how implicit and strategy-based motor adaption interact.

## INTRODUCTION

Accurate and precise perception of limb position is essential for movement control (Sanes et al., 1984). In recent years, the role of position sense for the flexibility of movements has attracted considerable attention, specifically its role for implicit motor adaptation (Tsay, Kim, et al., 2022). Implicit adaptation is an automatic, non-voluntary process (Mazzoni & Krakauer, 2006; Miyamoto et al., 2020), as opposed to explicit or strategy-based motor adaptation, which involves deliberation and cognitive effort (McDougle & Taylor, 2019; Taylor et al., 2014). Recent studies have linked implicit adaptation of arm movements to the precision with which people localize their hand (Henriques & T Hart, 2023; Tsay et al., 2021). Specifically, the asymptote of implicit adaptation, i.e., the maximum possible adaptation, depends on the precision, or inverse variability, of proprioception. However, previous studies have assumed that this precision remains static across learning. Here, we examined whether the variability of perceived hand position, as a proxy for precision, actually changes during motor adaptation.

Humans receive information about the position of their limbs from several sources of information. During reaching movements, signals from vision and proprioception are combined to form a coherent perceptual estimate of hand position (Van Beers et al., 1999). Any discrepancy in information between sensory modalities is reduced by two concurrent processes. These are proprioceptive recalibration, and (partial) multisensory integration (Block & Bastian, 2011). Proprioceptive recalibration results in a gradual shift of the perceived hand position towards visual feedback, which temporarily persists even when visual feedback is removed (Cressman & Henriques, 2009; Harris, 1963; Redding & Wallace, 1996). Proprioceptive recalibration has consequences for subsequent movements. Due to the shift in perceived hand position, the hand is initially perceived to miss the target in the direction of the visual feedback, even when it lands perfectly on target. This error in perceived hand position is thought to drive the gradual adaptation of subsequent movements in the direction opposite to the discrepant visual feedback, bringing visual feedback closer to the target (Henriques & Cressman, 2012; Tsay, Kim, et al., 2022). According to this view, implicit adaptation is therefore partly driven by proprioceptive recalibration.

However, in addition to recalibration, multisensory integration also occurs in the context of visuomotor adaptation (Rand & Heuer, 2020). Multisensory integration is a transient process that moves sensory estimates towards each other (Bosen et al., 2018; Ghahramani et al., 1997). The size of this shift for each modality depends on the precision, or inverse variability, with which sensory input from each modality is perceived. Less precise information is shifted more strongly (Debats et al., 2017; Ernst & Banks, 2002; Rand & Heuer, 2016). Therefore, suppressing proprioceptive information, and thereby increasing proprioceptive variability, may help reduce visuo-proprioceptive conflict (Limanowski, 2022).

Following the idea that implicit motor adaptation is driven by the discrepancy between perceived and desired limb position, lower proprioceptive precision, and, therefore, a larger shift of perceived limb position towards visual feedback, should result in stronger motor adaptation. Indeed, patients with loss of proprioceptive feedback move more accurately in some tasks with altered visuomotor contingencies than healthy controls (Lajoie et al., 1992). Besides its influence on the asymptote of implicit adaptation, proprioceptive precision may thus also influence the learning rate. It is therefore important to understand whether proprioceptive precision remains static, or changes during adaptation.

One reason why proprioceptive precision may change early during adaptation is attention. Attention modulates the neuronal gain in sensory areas (Limanowski & Friston, 2020). In a multisensory environment, such as during visuomotor adaptation, attention can increase perceptual precision of certain sensory modalities over others (Driver & Spence, 1998). During visuomotor adaptation, a shift in attention from proprioception to vision may be particularly pronounced when a cognitive strategy is used to compensate for the visual error. Strategy-based learning, which has been linked to eye movements towards the aiming position as an indicator of visuo-spatial attention (de Brouwer et al., 2018; Rand & Rentsch, 2015), may attenuate proprioceptive information, thereby reduce the precision of hand localization, and thus enhance implicit adaptation, as described above. This reduction in proprioceptive precision by intermodal attention is expected early during motor adaptation, when cognitive strategies influence adaptation most strongly (Taylor et al., 2014).

Hand localization reports represent a combination of information from various sources, including vision, efferent information, and proprioception. Imprecise proprioception should result in less precise, i.e., more variable, hand localization reports. We thus investigated whether the variability in hand localization increases at the beginning of visuomotor adaptation, and how such an increase is related to the use of a cognitive strategy.

Our experimental design alternated short movement cycles of visuomotor adaptation and washout, allowing us to repeatedly tap into an early stage of motor learning, and estimate proprioceptive precision by computing variance in perceived hand position across repetitions. To ensure robustness of our results, we employed two complementary methods for hand localization. We assumed that the strategy-based component of visuomotor adaptation, which includes shifting attention to vision over proprioception, attenuates proprioceptive precision. To test this, we examined hand localization during a combination of implicit and strategy-based learning (experiment 1), or under conditions of purely implicit learning (experiment 2). In a third experiment, we manipulated attention directly. Our results confirm that hand localization becomes less precise during early motor adaptation. We find that this decrease in precision occurs rapidly once a visuomotor rotation is introduced, and that it is most pronounced in the presence of a re-aiming strategy (experiment 1), and reduced when re-aiming is abolished (experiment 2), and when attention can be fully allocated to the hand (experiment 3).

## MATERIALS AND METHODS

### Subjects

Across the three experiments, 92 healthy, young volunteers took part. 30 participated in experiment 1 (10 female, average age 26 years, range 20-33 years), 33 in experiment 2 (11 female, average age 25 years, range 20-31 years), and 29 in experiment 3 (13 female, average age 26 years, range 22-33 years). These cohort sizes were determined based on extensive pilot testing. Participants were recruited via local participant databases and from staff and students of Otto-von-Guericke University and the Leibniz Institute for Neurobiology. They received a monetary reimbursement (8 Euros/hour). All participants were right hand dominant, verified by the Edinburgh handedness test, and had normal or corrected-to-normal vision. They gave written informed consent to the study protocol in advance to the experiment. The study was approved by the ethics committee of the university clinic Magdeburg, and conducted in accordance to the Declaration of Helsinki.

### Apparatus

Participants produced arm movements holding on to a two-link robotic manipulandum (Kinarm endpoint lab). The apparatus consisted of three levels. At the bottom level, participants held the handle of a robotic arm with each hand. Vision of their hands and arms was occluded by a semi-silvered mirror at the middle level, and a block cloth draped over their shoulders and arms. The mirror provided visual feedback from an LCD monitor (LG47LD452C, LG Electronics, 47 inch, 1,920×1,080 pixel resolution) mounted above at the top level with the display facing down. Distance between monitor and mirror was equal to distance between mirror and handles of the robotic arms, thus creating the illusion that visual feedback was in the same plane as the movement. Participants were seated in a comfortable chair with their forehead resting against a soft leather patch at the height of the top level. Kinematic data were recorded at a sampling rate of 1000 Hz, and with a spatial resolution of 0.1 mm. Occasionally, participants used a foot pedal for additional responses. The experiment was conducted in a dimly lit and quiet room.

### General experimental design

The goal of this study was to examine the precision of hand localization during the early stage of adaptation to a visuomotor rotation. Perceptual precision can be estimated from the variability of psychophysical reports, the computation of which requires a large number of repetitions. Our interest, however, was in the first few trials of adaptation, when strategy-based learning is most pronounced (Taylor et al., 2014). To capture the expected transient effects on perception, and nevertheless have sufficient repetitions for computing variability, we developed a new paradigm that required participants to repeatedly adapt and de-adapt for short cycles of few movements each. In each cycle, they reported their perceived hand position once, following the last movement of that cycle (as shown **Fig. 1a**).

**Fig. 1:**
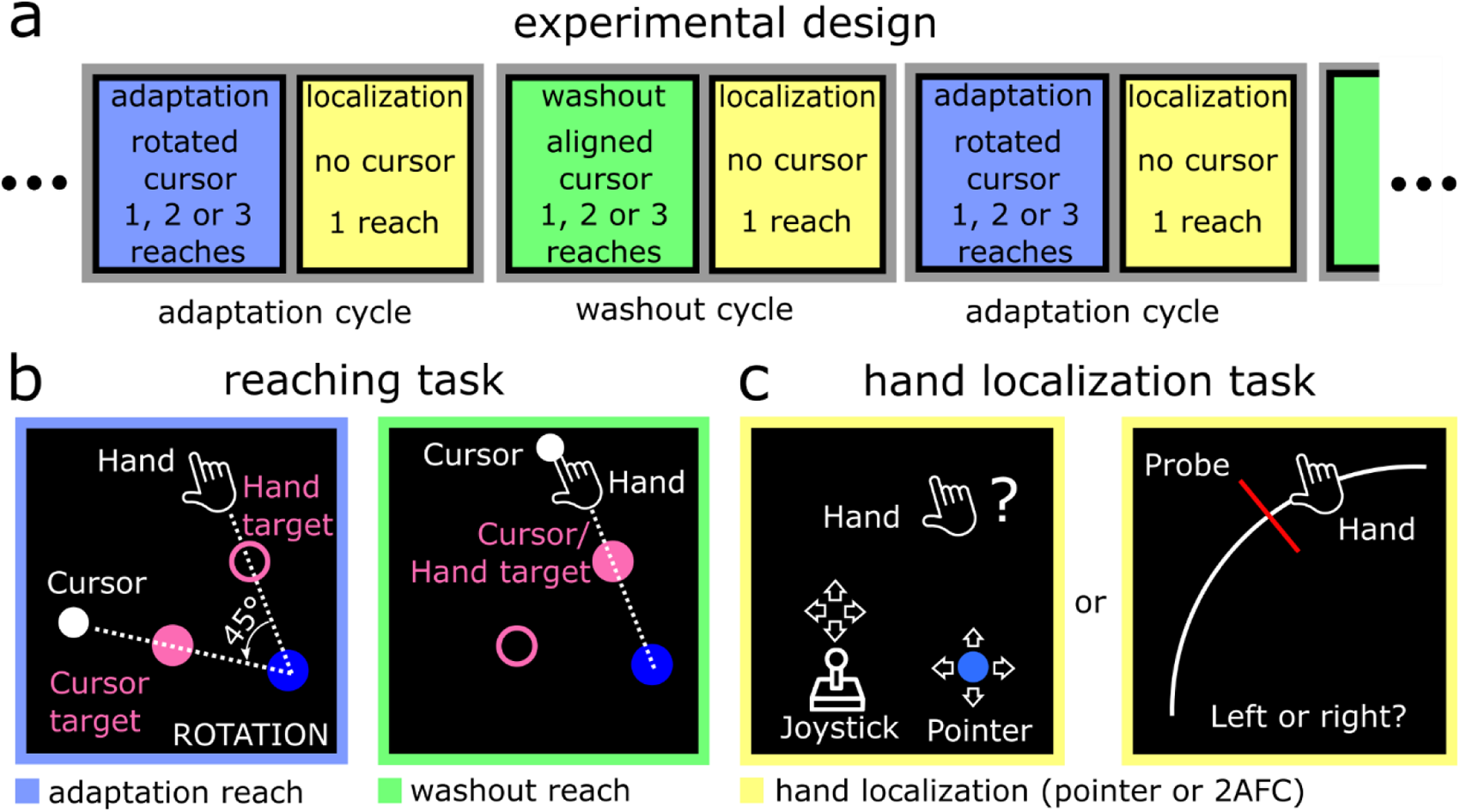
Experimental methods. (a) Schematic display of the task structure, in which short adaptation and washout cycles repeatedly alternated. Each cycle consisted of one, two, or three consecutive reaches, which were performed with 45° rotated (adaptation cycles, blue) or aligned (washout cycles, green) cursor. Each cycle ended with one reach that was performed without visual feedback, after which participants were asked to report their perceived hand position (yellow). (b) Participants had to move from a starting position (blue circle) through a target. In experiments 1 and 3 two targets were displayed (pink ring and dot). During adaptation cycles of experiments 1 and 3 (left), participants were instructed to use a compensatory movement strategy (“move through the hand target”) in order to make the cursor slice through the cursor target. In washout cycles (right), the cursor and hand target were identical, as cursor and hand position were congruent. In experiment 2, only one target was displayed, and participants were instructed to aim directly at the target during both adaptation and washout cycles (not shown here). (c) Participants reported their hand position at the end of the reach via one of two methods. They either moved a visual pointer to the exact location at which they perceived their right hand, using their left hand in a joystick fashion (left), or judged their right hand position to be left or right relative to a visual probe stimulus (right).

### Reaching task

Participants produced centre-out reaching movements with their dominant right arm in a horizontal plane, “slicing” through a visual target. Before each reach, the participants’ right hand was passively moved by the robotic manipulandum to the start position (blue circle with radius of 0.5 cm, 15 cm to the right from body midline, 15 cm from the front edge of the table. After keeping their hand inside the home position for 500 ms, the targets appeared at a distance of 10 cm from the home position. In experiments 1 and 3, two targets were displayed, a cursor target (pink unfilled circle in **Fig. 1b**, radius of 0.5 cm) and a hand target (pink filled circle, radius of 0.5 cm). In experiment 2, a single target (pink filled circle, radius of 0.5 cm) was displayed. Participants were instructed to produce rapid, straight movements and to stop shortly behind the target. There was no time limit for the initiation of the movement. However, once the movement was initiated, participants had to cross the target radius within 300 ms. Movement onset was defined as the point in time when a movement exceeded a velocity threshold of 5 cm/s, and a radial distance from the home position of at least 0.5 cm. If the time to cross the target radius exceeded 300 ms, they saw an error message on the display (“TOO SLOW”) and heard an error sound. After detection of movement offset (defined as velocity falling below a threshold of 5 cm/s), their hand was locked in endpoint position for 500 ms, and the trial ended.

Participants saw a cursor (white dot of 0.3 cm radius) as online visual feedback of the position of their right hand during active hand movements. The cursor was not shown during passive return movements. During adaptation cycles (blue-outlined panel in **Fig. 1b**), cursor movement was rotated 45° relative to the actual movement trajectory, while cursor and actual hand position were congruent during washout cycles (green outlined panel in **Fig. 1b**). Depending on the experiment, the rotation was either consistently 45° in clockwise direction, or clockwise (CW) or counterclockwise (CCW) in different cycles. Adaptation and washout cycles could consist of two, three, or four consecutive reaches, determined pseudo-randomly (equal number of cycles of each length). The number of reaches per cycle was varied to reduce predictability of the localization task, which concluded each cycle.

### Hand localization task

A single hand localization trial, as illustrated in the yellow-outline panels in **Fig. 1c**, concluded each cycle. These trials began much like the reach trials described above except that, upon movement onset, all visible stimuli disappeared, including the cursor and targets. In experiment 3, the cursor was not shown during hand localization at all. In all experiments, the right hand was held in the endpoint position by the robotic manipulandum until participants indicated their perceived hand position, and participants were instructed not to try to move the hand while it was locked in place. Throughout the experiments, we used one of two different hand localization methods, as follows.

### Pointer method

For hand localizations using the pointer method (the left yellow-outline panel in **Fig. 1c**), once the hand was locked in its endpoint position, a pointer (blue dot, radius 0.3 cm) appeared in a random location along an invisible, horizontal line in front of the participant (ranging from 5 cm to 21 cm to the right of the body midline, 10 cm from the front edge of the table). Participants had to move the pointer to the position where they perceived their right hand. They used their left hand to control the pointer by applying isometric force to the left robotic arm’s handle, which functioned similarly to a joystick. Once participants had aligned the pointer with their perceived right hand position, they had to press a pedal with their right foot to submit the pointer position as their psychophysical report. For each reported hand position, a graded performance score was calculated based on the Euclidean distance between the pointer and the actual hand position. Scores within a series of 36 adaptation and washout cycles (forming one experimental block) were summed to a total performance score, which was displayed at the end of each block to motivate accurate hand position reports.

### Two-alternative forced choice (2AFC) method

For hand localizations using the 2AFC localization method (right yellow-outlined panel in **Fig. 1c**), participants had to judge the position of a probe stimulus relative to the position of their right hand. Upon movement offset of the unseen hand, a white circular arc was displayed, spanning 140° (from 65° to 205° in polar coordinates), at a radius equal to movement distance. The actual right hand endpoint position was somewhere along the arc. A red tick mark was displayed on the arc at an angular offset of ±3.33°, ±10°, or ±20° relative the actual hand position (in pseudo-random order). Participants had to report if the probe was presented to the left or right relative to their right hand by pressing either a left or right foot pedal with their corresponding foot. If they answered correctly, a point was added to their overall performance score, which was not revealed to participants until the end of each experimental block.

### Experiment 1 – Re-aiming task

The goal of the first experiment was to examine if precision of perceived hand position decreases during the early stage of visuomotor adaptation. Specifically, participants performed the reaching task with short alternating cycles of adaptation and washout, as described above. During adaptation cycles, cursor movement was rotated 45° clockwise or counter-clockwise, creating a visuo-proprioceptive conflict that induces implicit motor learning. To induce strategy use, two targets were displayed (20° and 65° CCW relative the midline, ±7.5° jitter between cycles). Participants were instructed to move the cursor (white dot in **Fig. 1b**) through the cursor target (filled pink circle) by aiming for the hand target (unfilled pink circle) during adaptation cycles, i.e., to use an aiming strategy (see blue-outline panel **Fig. 1b**). As the angle between both targets corresponded exactly to the angle of visuomotor rotation, the instructed strategy could perfectly compensate for the rotation (Mazzoni & Krakauer, 2006). During washout cycles (green-outline panel **Fig. 1b**), cursor and hand position were aligned, and participants were instructed to move the cursor through the filled circle, while the unfilled circle was task-irrelevant. Visuomotor rotation direction, and the location of the filled and the unfilled circle, were reversed from one adaptation cycle to the next, so that participants needed to move to different targets in consecutive adaptation cycles. Thus, a rotation in CW rotation was associated with the instruction to aim for the target at 65° (bringing the cursor to the target at 20°), while a rotation in CCW rotation was associated with the instruction to aim for the target at 20° (bringing the cursor to the target at 65°). The required movement direction of washout cycles was matched with the previous adaptation cycle. A text below the start position indicated whether subjects were in a rotation cycle (“ROTATION”) so they could apply the instructed re-aiming strategy. During washout cycles, no text was displayed. Additionally, participants were informed that cycle types (adaptation or washout) would switch after each localization. Thus, participants knew from the beginning whether they were entering a rotation or washout cycle, and, thus, when to use the strategy, and when to stop using the strategy.

Participants were assigned to two groups, who indicated perceived hand position using different hand localization methods (pointer group and 2AFC group, each N=15). Both groups performed multiple blocks of the task, with each block consisting of 18 adaptation cycles alternating with 18 washout cycles (108 reaches and 36 localizations in total per block, circa 12 minutes per block). Between blocks, participants took a short break (2 minutes). The pointer group performed one baseline block, consisting of 36 cycles with aligned cursor movement, followed by six blocks of alternating adaptation and washout cycles. The number of blocks were chosen to allow for sufficient trial numbers to compute the shift and variability of hand localization for the second, third, and fourth reach in a cycle separately. Given that we observed no significant differences in localization variability between baseline and washout, we did not include a baseline block for the 2AFC group. Instead, the 2AFC group completed seven blocks of alternating adaptation and washout.

### Experiment 2 – Aim-direct task

In the second experiment, we tested if the reduction of hand localization precision during early visuomotor adaptation observed in experiment 1 depended on the use of a re-aiming strategy. Participants were instructed to ignore the visuomotor rotation during adaptation cycles, and aim directly at the target. This is a widely-used method to exclude strategic aiming (Hadjiosif & Krakauer, 2020; Maresch et al., 2021). As visual landmarks can promote the use of compensatory movement strategies (Taylor & Ivry, 2011) we presented only one target (at 45° relative to midline). Since there was only one target in this task, target jitter was expanded to ±15° to ensure a broad range of movements. Furthermore, as the alternating direction of visuomotor rotation may interfere with motor learning, visuomotor rotation was limited to CCW direction. Again, participants were assigned to two groups, who reported perceived hand positions using different hand localization methods (pointer group, N=15 or 2AFC group, N=18). As in experiment 1, the experiment consisted of multiple blocks, each containing 18 adaptation and washout cycles in alternating order. The pointer group performed six such blocks, while the 2AFC group completed seven blocks (three participants of the 2AFC group completed only six blocks).

### Experiment 3 – Cued localization

To investigate the influence of attention on the reduction of hand localization precision, we conducted a third experiment. During reaching, participants performed a re-aiming task as in the first experiment (see **Fig. 1b**). The visuomotor rotation was 45° in CCW direction. The hand target was always at 20° (±15° target jitter) relative to midline (cursor target 45° relative to hand target). In contrast to experiments 1 and 2, where the exact time point of a localization was unknown, participants in experiment 3 were explicitly informed beforehand whether a reach would be followed by the hand localization task. This was done by presenting a large warning symbol along with a text (“WHERE?”) on the screen from the beginning of localization trials onwards. In localization trials, participants could therefore fully focus attention on the position of their hand during the reaching movement. In one group, two targets were presented at the beginning of localization trials (up to the time of movement onset; 2-target group, N=15), and participants had to move their unseen hand through the hand target. In the other group, localization trials had only a single target. Given that all other trials had two targets, this additionally signalled the upcoming localization trial, and further minimised any attention away from the hand (1-target group, N=14). In both groups, perceived hand position was reported using the pointer method. Given that we did not observe differences in hand localization between the second, third, and fourth reach in a cycle in experiments 1 and 2, we analysed localization across different reach-numbers together in experiment 3, and could therefore reduce the number of hand localization trials. Participants performed three experimental blocks, as described above, resulting in 54 hand position reports for adaptation and washout, respectively.

### Data analysis

Data were analysed in MATLAB 2020b (The Mathworks Inc.). We recorded hand position throughout the experiment, and computed movement direction at maximum velocity, movement extent, and movement curvature offline based on hand position data for the outward movement. For experiment 1, we defined movement direction as positive in the direction of expected adaptation, i.e. in CW direction for the target associated with CCW rotation, and in CCW direction for the target associated with CW rotation. For experiments 2 and 3, movement direction was always defined to be positive in the CW direction, as rotation direction was always CCW. Movement extent corresponded to the radial distance from the home position to movement endpoint, i.e., the hand position at movement offset. Movement curvature was computed via the linearity index (LI), which was defined as the maximum perpendicular distance between the movement trajectory and a virtual line connecting home position and movement endpoint relative to movement extent (Atkeson & Hollerbach, 1985).

Perceptual reports obtained via the pointer method provided an angular error, defined as the angle between reported and actual hand position, both relative to the home position, as well as a radial error, defined as the difference in radial distance of the reported vs. actual hand position. The bias in perceived hand position was computed by averaging angular errors across localizations (after correcting for direction of rotation in experiment 1 by reversing the sign of angular errors in cycles with a CW rotation). The variability of hand localization reports (inverse precision) was computed using the inter-quartile-range (IQR) of angular errors (after de-meaning data for each target in experiment 1 separately). We chose IQR as a measure of variability due to its robustness to outliers.

Perceptual reports obtained via the 2AFC method were used to compute psychometric curves at the single-subject level. This was achieved by fitting a generalized linear regression model of the reported hand positions relative to the probe stimulus offset, for each cycle type (adaptation, washout) separately, using the glmfit.m function and a logit link function. Hand position reports were assumed to be binomially distributed. From the psychometric curves, biases in perceived hand position were computed as the point of subjective equivalence (PSE), i.e. as the (interpolated) probe stimulus offset for which subjects reported that the probe stimulus was “left” of their actual hand position on 50% of trials. Variability of hand localization reports was computed as the just noticeable difference (JND), i.e., the difference between the (interpolated) probe stimulus offsets for which subjects reported that the probe stimulus was “left” of their actual hand position in 75 vs. 25% of trials. For computation of the JND in experiment 1, curves for each target were first computed separately. The data were then combined by first subtracting the respective PSE from the probe stimulus offsets for each target, thereby effectively shifting the psychometric curves along the x-axis to align at x=0. A joint psychometric curve was then computed based on the x-shifted data points from both targets (as shown in **Fig. 3c**).

We rejected kinematic data of reaches that were too slow (movement duration>300 ms to reach target radius) or had a strong curvature, indicated by LI>0.2. In experiment 1, where two targets were presented, reaches towards the wrong target where excluded (movement direction<-30°). During pointer localization, participants occasionally pressed the foot pedal prematurely by accident. Therefore, hand position reports with high radial errors were rejected (defined as radial errors of hand position report that were less than half the actual movement extent, or radial errors that exceeded a threshold of 2 times the standard deviation of all radial errors).

The obtained data were analysed separately for adaptation and washout cycles. Kinematic data were furthermore binned depending on the reach-number in a cycle. For example, the adaptation aftereffect was computed as the mean movement direction during the first reach of all washout cycles. Similarly, pointer localization data in experiment 1 and 2 were binned depending on cycle length. Thus, we were able to compute biases and variability in perceived hand position for different time points, i.e., after two, three or four movements, during adaptation and washout. This allowed us to capture any short-term dynamics of perceived hand position.

### Statistics

Statistics were computed in MATLAB2020b and JASP 0.14.0.0 (JASP Team, 2020). We report mean values and standard deviation or, when a Shapiro-Wilk test indicated a violation of the assumption of normality, we report median values and inter-quartile-range. Mixed analysis of variance (ANOVA) was used to test for differences in movement kinematics between different conditions, and between participants of different groups. Perceptual variables obtained from pointer localizations were submitted to repeated ANOVAs. Perceptual variables from 2AFC localizations were analysed using paired-samples t-tests. When dependent variables were not distributed normally we used a corresponding non-parametric test. T-test were one-sided when we tested for effects previously reported in the literature, and two-sided when effects were not known a priori from the previous literature. We used mixed and repeated ANOVAs, as well as paired-samples t-tests for additional post-hoc analysis. Bayesian t-tests were used where the goal was to test for evidence in favour of the null hypothesis.

### Data availability statement

All raw data is available under doi: 10.5281/zenodo.8319018

## RESULTS

### Experiment 1 – Re-aiming task

In the first experiment, our primary hypothesis was that the precision of perceived hand position decreases during early adaptation to a visuomotor rotation. We expected adaptation to be evident in a gradual overcompensation of movement direction (Mazzoni & Krakauer, 2006), i.e., an increase in cursor error from the first to the fourth reach during adaptation, accompanied by a bias in hand localization towards the rotated visual feedback.

#### Adaptation of reaching movements

We found small cursor errors (angular error between cursor and cursor target) during adaptation cycles, indicating that participants successfully applied the instructed re-aiming strategy to compensate for the visuomotor rotation. Instead of the expected gradual overcompensation (Mazzoni & Krakauer, 2006), we found that cursor error was significantly greater than zero during the first reach of adaptation cycles, and afterwards decreased across subsequent adaptation reaches (as illustrated by blue circles from a1 to a4 in **Fig. 2a**). We performed a 2x4x2-mixed-effects ANOVA to test for effects of the within-subject factors Cycle Type (adaptation and washout) and Reach-number (i.e. 1^st^ reach, 2^nd^ reach, 3^rd^ reach, 4^th^ reach in a cycle) on cursor error. In addition, we included the between-subjects factor Group (pointer group and 2AFC group). We found a main effect for factors Cycle Type (F(1,28)=4.49, p=.04, *η*^2^=.06) and Reach-number (F(3,84)=39.98, p<.001, *η*^2^=.02), and an interaction between Cycle Type and Reach-number (F(3,84)=4.79, p=0.004, *η*^2^=.003). Post-hoc tests showed a significant decrease in cursor error from the first to the following reaches of washout (all t(29)>5.90, all p<.001 for 1^st^ vs. 2^nd^, 1^st^ vs. 3^rd^ and 1^st^ vs. 4^th^, Holm correction), as illustrated by green circles from w1 to w4 in **Fig. 2a**. Cursor error decreased also across adaptation cycles (t(29)=4.46 for 1^st^ vs. 3^rd^, t(29)=6.09 for 1^st^ vs 4^th^, both p<.001, Holm correction). Adaptation between both groups did not differ significantly (all p-values for interactions with the factor Group > 0.1). We therefore collapsed data across the two groups for further kinematic analysis.

While the decrease of cursor error during adaptation cycles is inconsistent with the overcompensation reported as evidence of implicit learning during instructed strategy use in previous studies (Mazzoni & Krakauer, 2006), we found aftereffects during washout cycles, which are considered a hallmark of implicit adaptation (see **Fig. 2a**; green). Furthermore, aftereffects increased with the number of immediately preceding rotated reaches (F(2,58)=22.17, p<0.001, *η*^2^=.43, **Fig. 2b**; t(29)=-3.19, p=.002, two vs. three preceding reaches; t(29)=-3.47, p=.002, three vs. four preceding reaches; Holm correction), providing evidence of gradual implicit learning throughout adaptation cycles.

Comparing the dynamics of cursor error during adaptation between early and late stages of the experiment (first vs. last 30 adaptation cycles), we found evidence that subjects did not simply use the instructed strategy, but adjusted it throughout the experiment via ongoing strategy-based learning. Early during the experiment, cursor error increased by 2.41°±3.41° from the first to the second reach of adaptation cycles (t(29)=3.77, p<.001, d=.69), consistent with gradual overcompensation. However, at later stages of the experiment, cursor error decreased by -1.02°±2.02° (t(29)=-2.76, p=.01, d=.50), consistent with the idea that subjects adjusted the instructed aiming strategy across the course of the experiment (see also **Supplementary Analysis S1**).

**Fig. 2:**
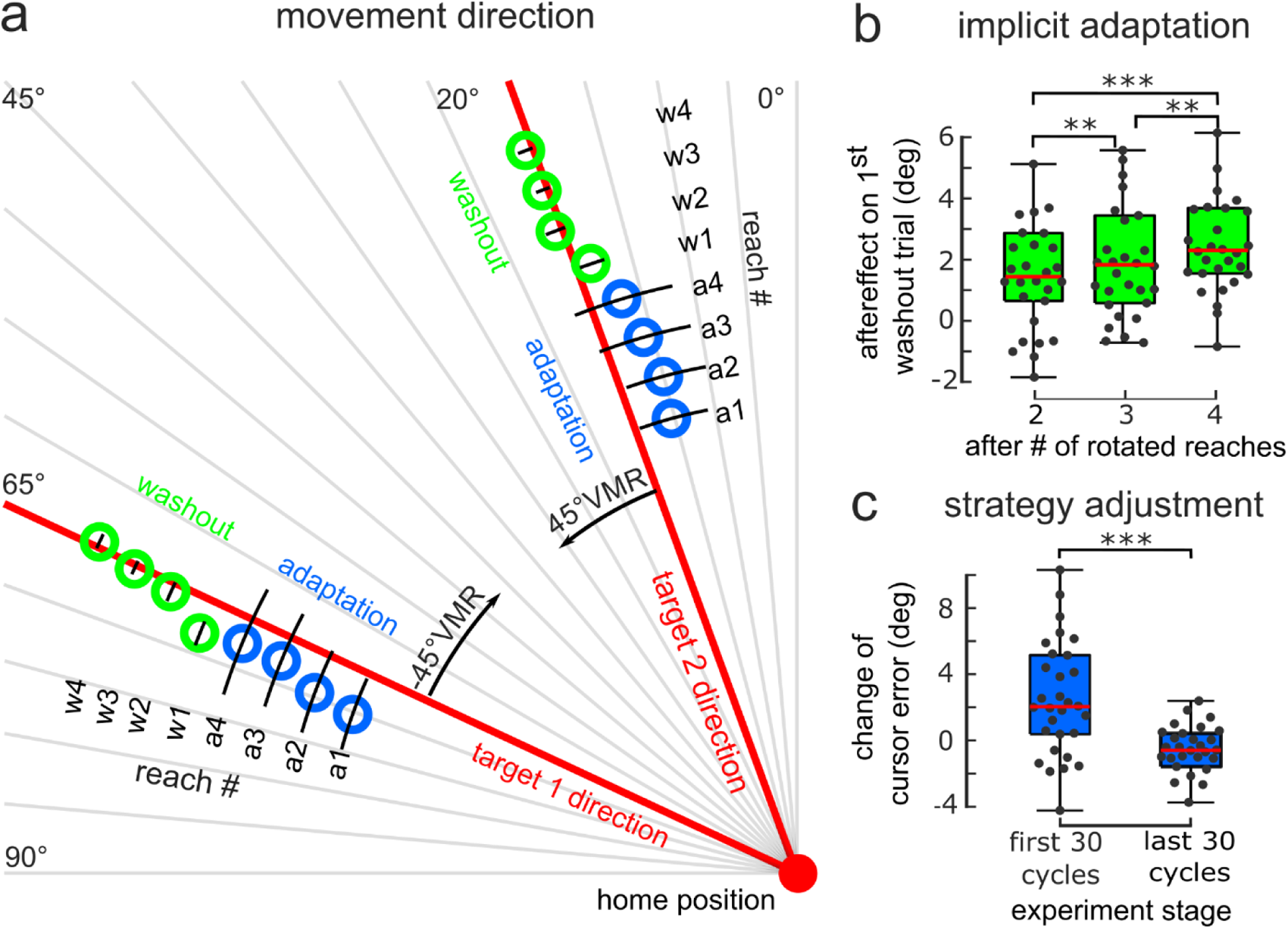
Adaptation of reaching movements in experiment 1. (a) Movement direction (blue and green rings, black lines indicate std, N=30, both groups) towards both targets (target direction indicated by red lines) were shifted in opposite direction of the rotated visual feedback (±45° VMR) throughout the reaches of adaptation cycles (“a”, numbers and radial extent from home position represent order of reaches, e.g. “a1” being the first reach of adaptation cycles, “a2” the second reach of adaptation cycles, etc.). The first reach of washout cycles (“w1”) showed an adaptation aftereffect, which decreased over consecutive movements (“w2”, etc.). (b) Implicit motor adaptation was evident as an increase in aftereffects (boxplots, dots represent different subjects, red line shows median) with the number of preceding reaches performed with rotated visual feedback. (c) Change of cursor error from 1^st^ to 2^nd^ reach in adaptation cycles was positive during the first 30 cycles of the experiment (left boxplot, dots represent different subjects, red line shows median) but negative during the last 30 cycles (right). Positive values reflect overcompensation. Negative values reflect adjustment of aiming strategy. **p<.01, ***p<.001

#### Changes in hand localization

As expected, we found that perceived hand position was biased towards the rotated cursor during adaptation cycles for both the pointer and the 2AFC group (**Fig. 3a**). For the pointer group, angular hand localization errors during adaptation and washout cycles were 1.96°±6.18° and 0.46°±3.05° (median±IQR), respectively (F(1,14)=4.63, p=.049, *η*^2^=.23; detailed statistics in **Supplementary Analysis S2**). In the 2AFC group^1^, localization bias was 1.69°±3.55°during adaptation, and -0.74°±4.38° during washout cycles (t(13)=2.27, p=0.02, d=.61, one sided; **Fig. 3d**).

Importantly, in both groups, we also found an increase in inter-quartile-range (IQR) of angular localization errors during adaptation cycles compared to washout cycles. For the pointer group, the IQR during adaptation cycles was 10.58°±4.98°, while IQR for washout cycles was 6.02°±1.83°, i.e., comparable to an IQR of 6.78°±2.48° during baseline. We conducted a 2x3-rANOVA with within-subject factors Cycle Type and Reach-number, and found an effect of Cycle Type (F(1,14)=15.67, p=.001, *η*^2^=.36, **Fig. 3b**). There was no significant effect of Reach-number (F(2,28)=.58, p=.57), nor an interaction between both factors (F(2,28)=.06, p=.95).

**Fig. 3:**
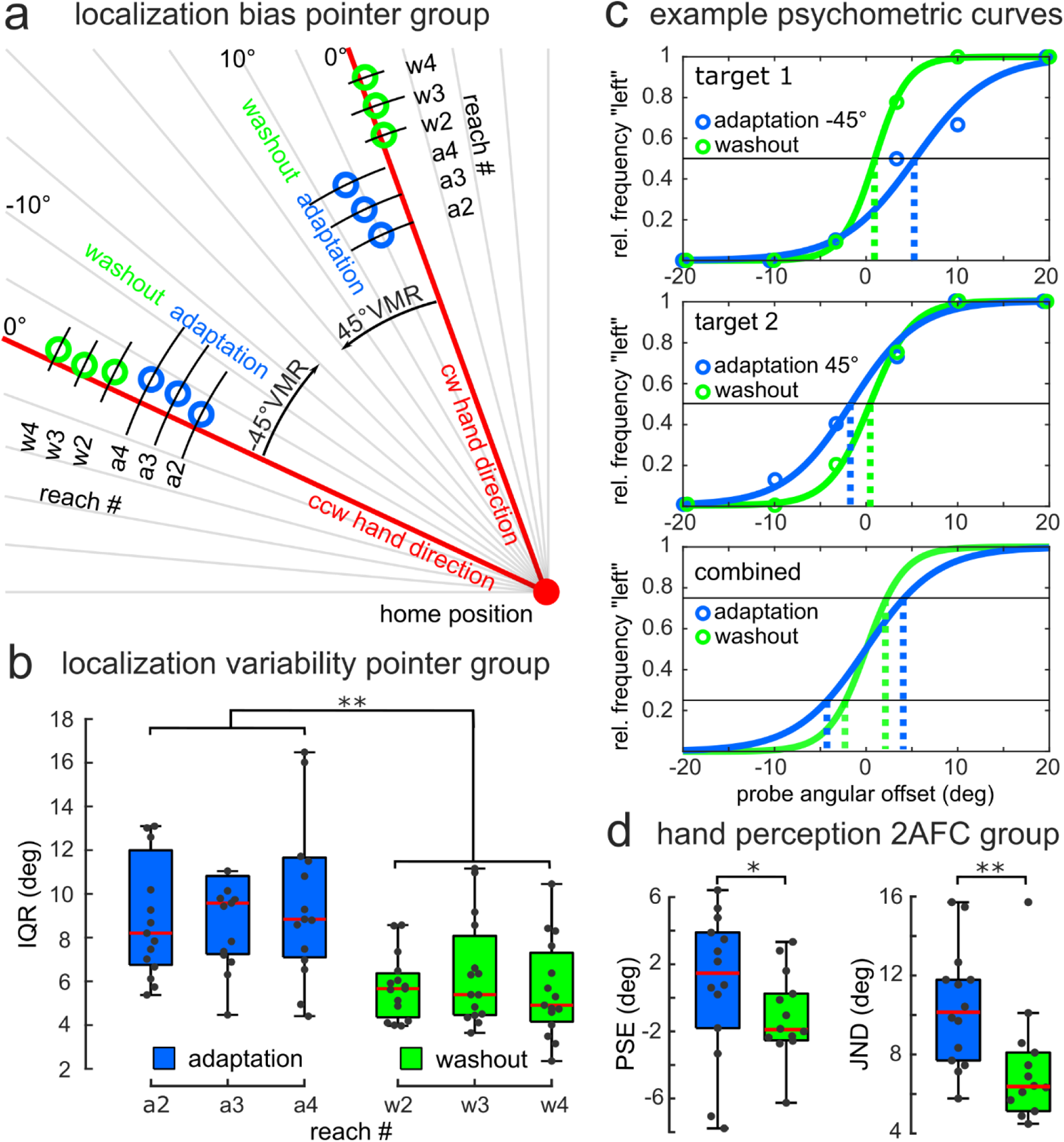
Changes in hand localization in experiment 1. (a) Median hand position reports for each reach during adaptation (blue rings; reaches a1-a4, black lines indicate IQR) and washout (green rings; reaches w1-w4). Target direction is indicated by the red lines. Hand localization was biased towards the rotated cursor (±45° VMR) in adaptation (“a”, blue) compared to washout cycles (“w”, green). Numbers and radial distance from the home position indicate the reach-number in a cycle at which participants reported their hand position, e.g. “a2” after the second movement in an adaptation cycle (the first reach never required localization. (b) Inter-quartile-range (IQR, boxplots, dots represent individual participants, red line shows median) of hand localization errors was increased during adaptation cycles, compared to washout cycles across all reach-numbers (e.g. “a2” the second movement in adaptation, etc.) (c) Example psychometric curves (solid lines) fitted to 2AFC reports (circles) for target 1 (upper panel) and target 2 (middle panel). For each participant in the 2AFC group, the point of subjective equivalence (PSE) was computed for both targets (dashed lines, note different direction of localization bias for different rotation directions), and subsequently subtracted from the respective probe angular offset values. The unbiased probes for both targets/ rotation directions were then combined to compute the just-noticeable-difference (JND, lower panel, interval between dashed lines). (d) PSEs were biased in adaptation, compared to washout cycles (left). JND was increased in adaptation compared to washout cycles (right). *p<.05, **p<.01; N=15 for panels a and b, and N=14 for panel d.

For the 2AFC group, the average JND were 10.41°±3.07° for adaptation, and 6.87°±3.45° for washout. A paired t-test in this group showed that the JND differed significantly between both cycle types (t(13)=3.99, p=0.002, d=1.07, **Fig. 3d**). Control analyses excluded the possibility that the observed difference in hand localization variability could be explained by a systematic drift in localization bias across the experiment, or by increased variability in hand positions during localization in adaptation cycles (see **Supplementary Analysis S3**)

### Experiment 2 – Aim-direct task

We next asked whether the observed increase in hand localization variability during adaptation was limited to conditions under which subjects employ re-aiming strategies, or exists more generally even when learning is purely implicit. In experiment 2, we thus instructed participants to ignore the rotation of the cursor relative to movement direction during adaptation cycles and aim directly at the target. Due to the instruction to move the hand through the target, we report the movement direction, rather than the cursor error as in experiment 1.

#### Adaptation of reaching movements

We performed a 2x4x2-mixed-ANOVA with the within-subject factors Cycle Type (adaptation and washout) and Reach-number (i.e. 1^st^ reach, 2^nd^ reach, 3^rd^ reach, 4^th^ reach in a cycle). Additionally, we included the between-subjects factor Group (pointer group and 2AFC group) to confirm that the localization method had no effect on movement adaptation. We found a main effect for factors Cycle Type (F(1,31)=65.93, p<.001, *η*^2^=.23, **Fig. 4a**), and an interaction between Cycle Type and Reach-number (F(3,93)=76.23, p<.001, *η*^2^=.11). Movement direction gradually shifted in CCW direction from the first to the fourth movement during adaptation cycles (all t(28)<-7.84, all p<.001 for 1^st^ vs. 2^nd^, 1^st^ vs. 3^rd^, 1^st^ vs. 4^th^, t(28)=-2.85, p=.03 for 2^nd^ vs. 3^rd^, Holm correction), and reverted during washout cycles (all t(28)>7.74, all p<.001 for 1^st^ vs. 2^nd^, 1^st^ vs. 3^rd^, 1^st^ vs. 4^th^, Holm correction). Adaptation between both groups did not differ significantly (all p-values for interactions involving Group > 0.6). Changes in movement direction were much smaller than the visuomotor rotation, so that the cursor during adaptation cycles obviously missed the target, indicating that participants did not apply a compensatory movement strategy during adaptation cycles, and that motor learning was implicit.

#### Changes in hand localization

As in experiment 1, we found a bias in hand localization in the direction of the rotated cursor during adaptation cycles for both groups^2^. Examining angular localization errors as a function of Reach-number in a cycle in the pointer group, we found that the perceived hand position was biased towards the cursor after a single movement with rotated visual feedback (see **Fig. 4b**, for statistical analysis see **Supplementary Analysis S2**).

To test if IQR of angular localization errors was again higher during adaptation, compared to washout, we performed a 2x3-rANOVA for the pointer group with within-subject factors Cycle Type (adaptation or washout) and Reach-number (i.e. 2^nd^, 3^rd^, or 4^th^ reach in a cycle). This revealed a significant effect of Cycle Type, with higher hand localization IQR during adaptation (8.83°±1.93°), compared to washout (7.50°±2.05°; F(1,14)=6.34, p=0.03, *η*^2^=.15, **Fig. 4c**). We did not find a significant effect of Reach-number (F(2,28)=1.09, p=.35), nor an interaction between both factors (F(2,28)=.86, p=.43).

However, for the 2AFC group, we did not find a significant difference between angular localization error JND during adaptation and washout cycles (t(13)=.02, p=.99). We performed an additional Bayesian t-test, and found moderate evidence in favour of the null hypothesis, i.e., that JND in both cycle types were equal (BF_10_=0.27).To explain these discrepant findings between the pointer group and the 2AFC group, we considered a potential role of differences in the role of the target during hand localization in an additional post-hoc analysis (see **Supplementary Analysis S4**).

**Fig. 4:**
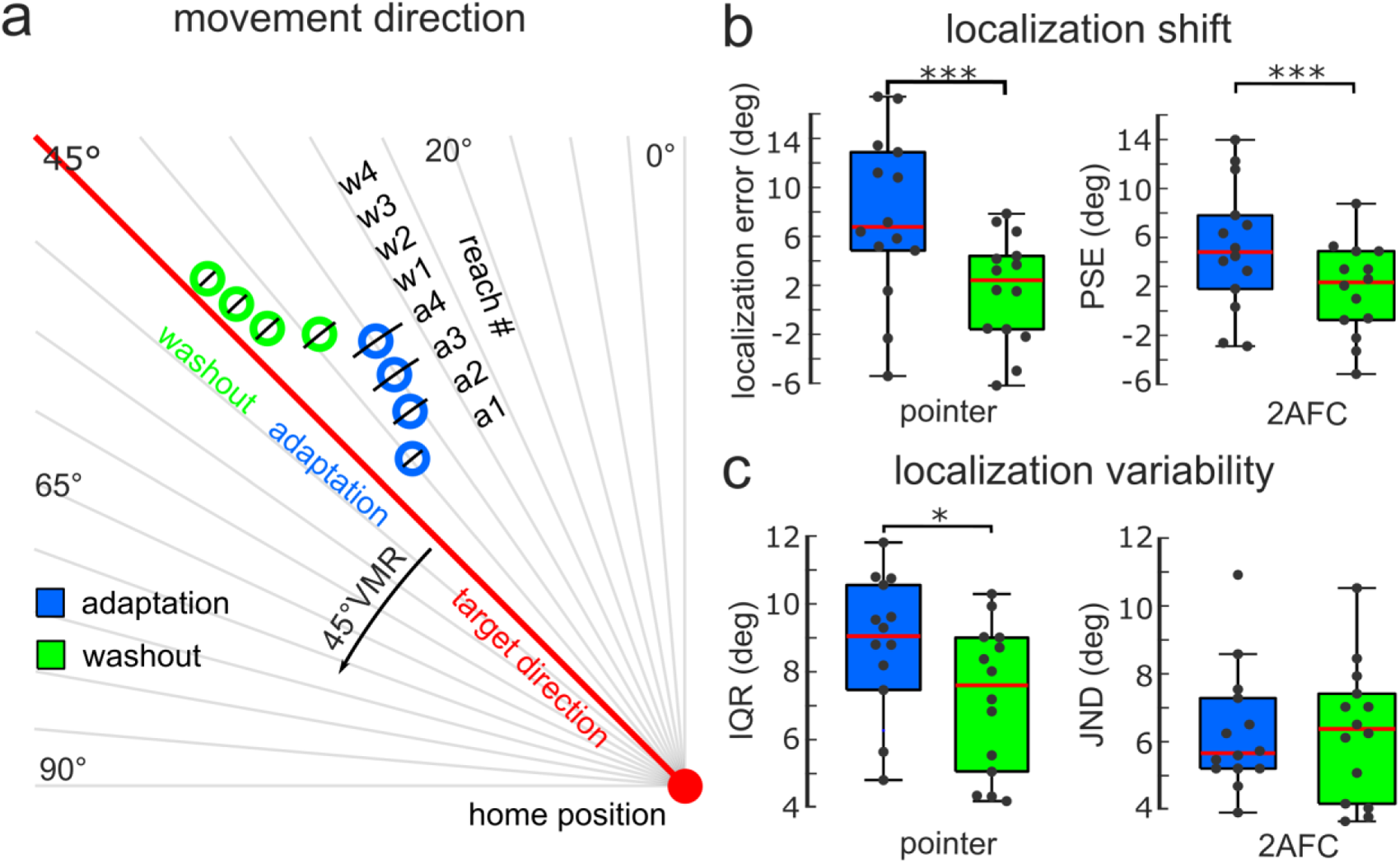
Kinematic and perceptual results experiment in 2. (a) Mean movement direction for each reach during adaptation (blue rings; reaches a1-a4) and washout (green rings; reaches w1-w4; N=33, across groups). The black lines indicate standard deviation of movement direction. Target direction is indicated by the red line. Movement direction was shifted in the direction opposite to the rotated visual feedback (±45° VMR) for all reaches during adaptation cycles (“a1” being the first reach of adaptation cycles, “a2” the second reach of adaptation cycles, etc.). The first reach of washout cycles (“w1”) showed an adaptation aftereffect, which decreased over consecutive movements (“w2”, etc.). The radial distance from the home position at which blue and green rings are positioned does not represent movement extent, but displays the time-course of consecutive reaches (a1 to a4, then w1 to w4). (b) Hand position reports (boxplots, dots represent single participants, red line shows median) were shifted in the direction of visual feedback during adaptation (blue) compared to washout cycles (green) for pointer (left, N=14) and 2AFC (right, N=14) groups. (c) Variability increased in adaptation compared to washout cycles for the pointer group (left). However, we did not find a difference between the two cycle types for the 2AFC group (right). *p<.05, ***p<.001

### Experiment 3 – Cued localization

In experiment 3, we tested whether the decrease in hand localization precision observed in experiment 1 persisted even when participants could fully focus attention on the position of their hand during localization. To this end, participants completed a close variant of experiment 1; however, hand localizations were pre-cued, and therefore predictable. As we found evidence in experiment 2 that the 2AFC method may emphasize an influence of remembered target location on hand localization, experiment 3 employed the pointer method.

#### Movement adaptation

We performed a 2x4x2-mixed-ANOVA with within-subject factors Cycle Type (adaptation or washout) and Reach-number (i.e. 1^st^ reach, 2^nd^ reach, 3^rd^ reach, 4^th^ reach in a cycle), and between-subject factor Group (2-target or 1-target). We found a significant effect of Cycle Type (F(1,27)=13.41, p=0.001, *η*^2^=.08, **Fig. 5a**), and an interaction between Cycle Type and Reach-number (F(3,81)=43.64, p<0.001, *η*^2^=.07). The interaction was due to a gradual increase of cursor error from the first over subsequent reaches during adaptation cycles (1^st^ vs. 2^nd^, 1^st^ vs. 3^rd^, 1^st^ vs. 4^th^, all t(28)<-5.68, all p<.001, Holm correction, as illustrated by blue circles from a1 to a4 in **Fig. 5a**), and a decrease of cursor error during washout cycles (1^st^ vs. 2^nd^, 1^st^ vs. 3^rd^, 1^st^ vs. 4^th^, all t(29)>5.01, all p<.001, Holm correction, as illustrated by green circles from w1 to w4 in **Fig. 5a**). As in experiment 1, participants used a compensatory aiming strategy during adaptation in these experiments. Unlike experiment 1, experiment 3 did not reveal any adjustment of the aiming strategy across the experiment, consistent with the reduced number of blocks in experiment 3 (change of cursor error from 1^st^ to 2^nd^ reach in adaptation, first 30 cycles: 3.30°±4.05°, last 30 cycles: 2.97°±4.88°, t(27)=.47, p=.64, BF_01_=4.50). There was neither an effect of Group, nor an interaction of Group with either of the two within-subject factors (all p>.1).

#### Changes in hand localization

As in previous experiments, angular hand localization errors were biased towards the rotated cursor in adaptation cycles, compared to washout cycles (see **Fig. 5b**, for statistical analysis see **Supplementary Analysis S2**).

A 2x2-mixed-ANOVA of hand localization variability with within-subject factor Cycle Type, and between-subject factor Group, revealed a significant effect of Cycle Type (adaptation: 9.50°±3.28°, washout: 7.61°±3.52°, F(1,27)=18.83, p<0.001, *η*^2^=.07, **Fig. 5c**), but no effect of Group (F(1,27)=0.01, p=0.92), nor an interaction between Cycle Type and Group (F(1,27)=2.50, p=0.13). Consistent with results from experiment 1, hand localization reports were thus more variable in adaptation cycles, compared to washout cycles.

**Fig. 5:**
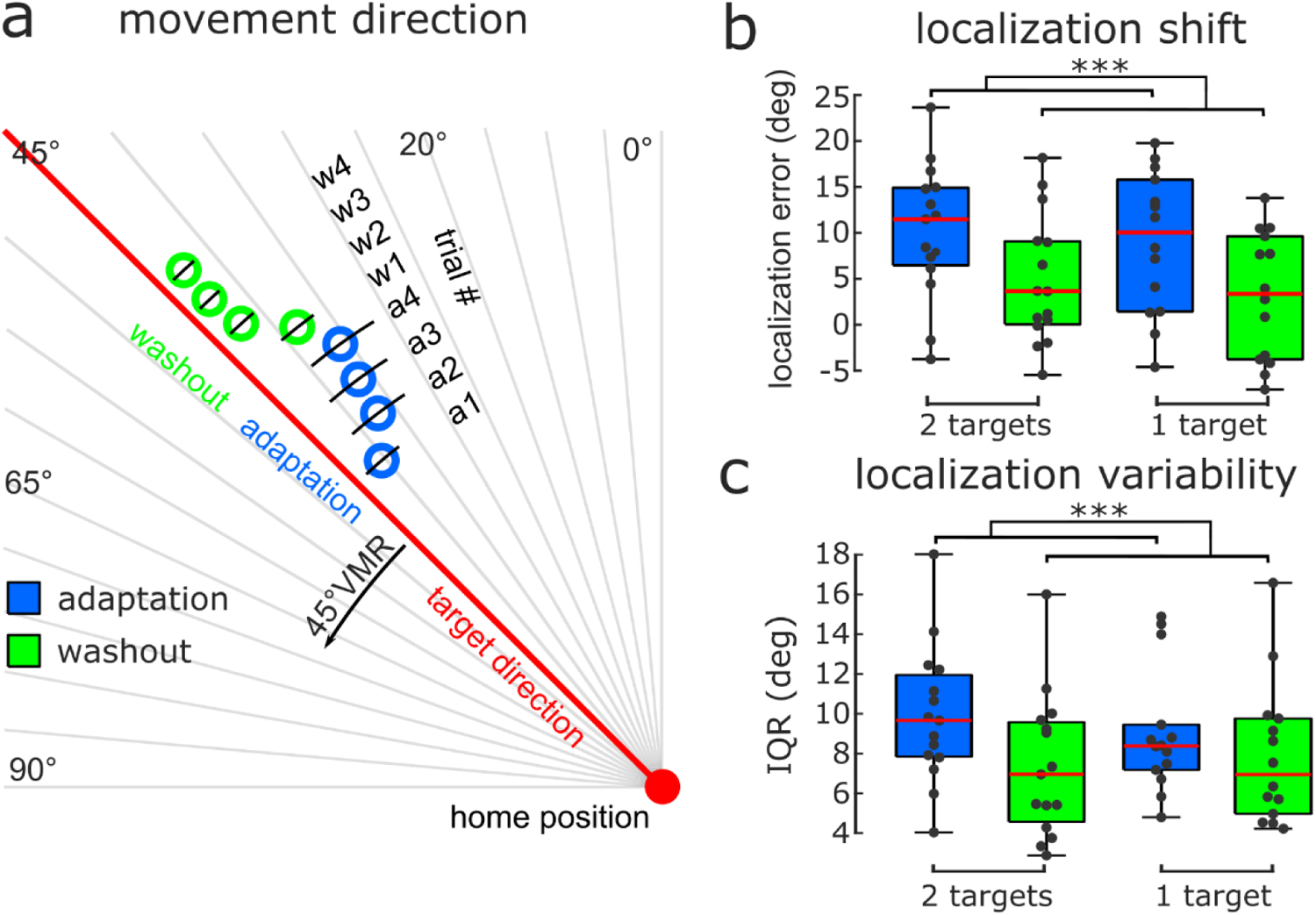
Kinematic and perceptual results in experiment 3. (a) Mean movement direction for each reach during adaptation (blue rings; reaches a1-a4, black lines indicate std) and washout (green rings; reaches w1-w4; N=28, across groups). Target direction is indicated by the red line. Movement direction was shifted in the direction opposite to the rotated visual feedback (45° VMR) for all reaches during adaptation cycles (“a1” being the first reach of adaptation cycles, “a2” the second reach of adaptation cycles, etc.). The first reach of washout cycles (“w1”) showed an adaptation aftereffect, which decreased over consecutive movements (“w2”, etc.). The radial distance from the home position at which blue and green rings are positioned does not represent movement extent, but displays the time-course of consecutive reaches (a1 to a4, then w1 to w4). (b) Hand position (boxplots, dots represent individual participants, red line shows median) reports were shifted in the direction of visual feedback during adaptation (blue) compared to washout cycles (green) for 2-target (left, N=14) and 1-target (right, N=14) groups. (c) Variability increased in adaptation compared to washout cycles for the 2-target (left) and 1-target groups (right). ***p<.001

## DISCUSSION

We report that the precision with which subjects can localize their hand changes during early motor adaptation. Our results show that precision of hand localization decreases rapidly once a visuomotor rotation is introduced in a reaching task. This decrease is observed when subjects use a compensatory aiming strategy (experiment 1), and persists even when subjects can fully allocate attention to hand position (experiment 3), and, at least under certain circumstances, when learning is purely implicit (experiment 2). Position sense in general, and its precision in particular, are increasingly recognized as important determinants of motor adaptation (Henriques & T Hart, 2023; Tsay, Kim, et al., 2022). Hence, the precision decrease observed during early adaptation cycles in our study may have important implications for our understanding of how movements are adapted.

We discuss how the observed dynamics in hand localization could relate to mental processes associated with implicit or strategy-based learning, outline potential consequences of changes in the precision of hand localization for motor learning, and finally discuss limitations of our study, together with potential future directions.

### Component processes of adaptation that may impact on localization precision

We observed the largest decrease in hand localization precision in experiment 1, when movements were governed both by strategy-based learning and implicit adaptation. Experiment 2 revealed that this decrease can persist in the absence of any re-aiming strategy, when adaptation is purely implicit, however, only under certain circumstances that depend on the reporting method. Furthermore, even for the pointer group in experiment 2, for whom we observed a significant increase in variability, this increase was significantly smaller than in experiment 1 (see **Supplementary Analysis S5**). This supports the idea that hand localization becomes less precise during adaptation in particular when cognitive strategies are involved.

Cognitive re-aiming strategies in visuomotor adaptation likely shift attention from the hand to the (rotated) visual feedback. Such intermodal attention results in a synaptic gain modulation that down-weights somatosensory information (Limanowski, 2022). We therefore propose that one reason why hand localization becomes less precise during adaptation is an intermodal shift of attention during strategy use.

Consistent with this idea, hand localization variability increased less in experiment 3, compared to experiment 1 (see **Supplementary Analysis S5**). As we have shown, participants in experiment 1 adjusted the instructed strategy throughout the experiment, in line with previous reports (e.g., Taylor & Ivry, 2011). It seems plausible that this continuous strategy adjustment caused uncertainty in the aiming location and involved careful monitoring of the relation between (rotated) visual feedback and cursor target, i.e., attention to vision, rather than proprioception. In the localization trials of experiment 3, on the other, participants could fully focus on hand localization, rather than shifting attention to vision, and aim directly at the visual target. This may explain why the increase in variability of hand localization was less pronounced in experiment 3.

Interestingly, a significant increase in variability nevertheless persisted in experiment 3. This indicates that the recent history of strategy use resulted in an enduring decrease of precision even in localization trials, when subjects could fully attended to hand position. The observed decrease in precision is therefore more than a short-lived result of momentary inattention. While the underlying mechanism interacts with attention, it is not attention itself.

Experiment 2 showed that the observed decrease in precision can be resistant to aiming directly at the target, depending on the reporting method. However, while participants were instructed to ignore the cursor, attention may nevertheless play an important role for the observed decrease in precision. Vision typically conveys essential information about limb positions, in particular in relation to the external world. It may be therefore difficult to ignore vision completely. For example, visual feedback that is invariant to movement direction (also referred to as error-clamped feedback) drives feedforward motor adaptation even though participants are made aware of the manipulation, and instructed to ignore the visual feedback (Kim et al., 2018; Morehead et al., 2016).

Besides attention, another factor that may contribute to the decrease in precision is the prediction error that a visuomotor rotation necessarily introduces. The perception of limb position does not solely rely on somatosensory brain areas; the cerebellum also plays a role in position sense during active movements (Weeks et al., 2017b, 2017a). Predictive computations of the cerebellum may enhance the precision of position sense (Bhanpuri et al., 2013). Similar to integration of multiple sensory modalities, combination of sensory and predictive information may depend on their respective uncertainty (Vaziri et al., 2006). However, when predictive models are updated to a change in visuomotor contingency, their estimation uncertainty is thought to increase due to the presence of prediction errors, which are most pronounced in the initial stage of motor adaptation (Tan et al., 2016). Therefore, another possible explanation is that the precision of hand localization decreased during early visuomotor learning in our study due to less precise predictive information. However, if prediction error played a major role in the observed decreasing in hand localization precision, we would have expected to observe a similar decrease also in washout cycles, in which participants experienced errors between predicted and perceived movement outcomes as well, evident in adaptation aftereffects (albeit likely of substantially smaller size). This was not the case, nor did we find a deterioration of perceptual precision between the baseline condition and washout cycles in experiment 1.

### Dynamics of perceptual changes during visuomotor adaptation

Previous work on the neural processing of somatosensory feedback during visuomotor adaptation suggests a suppression of proprioceptive information during the early period of learning (Bernier et al., 2009; Jones et al., 2001; Limanowski & Friston, 2020). However, these neurophysiological studies have left the question unanswered whether the precision of perception changes. The few psychophysical studies that have investigated variability of position sense in a context of visuomotor adaptation have done so at a late stage of adaptation, and found no decrease in precision (Cressman & Henriques, 2009; Tsay et al., 2021).

This may not be surprising, given that a decrease of precision is more likely during the early period of motor learning, when re-aiming strategies dominate learning. Indeed, we found a decrease that was already present after a single reaching movement with rotated visual feedback. Precision recovered immediately and completely during washout cycles. Importantly, hand localization trials were identical across both types of cycles. That is, the reaching movements during these trials were performed in the absence of visual feedback, both in adaptation and washout cycles. Precision of hand localization therefore did not depend on the immediate visual feedback in the current trial, but likely on the focus of attention directed by the task requirements during the present reach (experiment 1), and in recent trials (experiment 3). Furthermore, precision did not change across the three reaching movements of an adaptation cycle. However, we expect precision of hand localization to recover over the course of visuomotor adaptation, as previous studies reported that variability of proprioception during the late stage of adaptation is not different from baseline (Cressman & Henriques, 2009; Tsay et al., 2021). This could be because participants likely rely less on visual attention during the late stage of adaptation, when a successful re-aiming strategy has already been established (Reuter et al., 2015).

In addition to a decrease in hand localization precision, we observed that the visuomotor rotation introduced a systematic bias in localization, consistent with recalibration of proprioception. Cross-sensory recalibration has been regarded as a gradual process (Debats et al., 2023; Zbib et al., 2016). Here, we observed a notably fast localization bias, occurring even after a single reaching movement with rotated visual feedback. This finding confirms previous studies, which have reported similarly fast shifts for proprioception (Ruttle et al., 2018, 2021). Importantly, perceptual changes did not increase over time, speaking against the idea of a gradual change. Hand localization was performed in the absence of visual feedback and should thus reflect recalibration of proprioception, but not multisensory integration, as the latter requires simultaneous input from two sensory modalities.

### Potential consequences of a decrease in hand localization precision for motor learning

A recent model casts implicit motor adaptation as the result of an error between desired and sensed hand position, where the latter is derived from a proprioceptive estimate (Tsay, Kim, et al., 2022). During visuomotor rotation, proprioception is shifted towards visual feedback due to sensory recalibration and partial multisensory integration (Rand & Heuer, 2020). The rules of multisensory integration are governed by the precision of each sensory estimate, such that less precise estimates are shifted more strongly (Rand & Heuer, 2020). Reduced proprioceptive precision during visuomotor adaptation should thus result in a stronger shift of the estimated hand position towards visual feedback due to (partial) multisensory integration. As a consequence, following the idea that a shift in proprioception contributes to implicit adaptation, a reduction in proprioceptive precision during visuomotor adaptation should enhance implicit adaptation. Thus, the dynamic change of precision in hand localization observed here might act as a mechanism that controls the speed of adaptation.

We found that the precision of hand localization was lowest in experiment 1, when participants used a compensatory aiming strategy. Thus, we would expect strategic re-aiming to enhance implicit adaptation. This is because lower precision has been associated with stronger implicit motor adaptation (Henriques & T Hart, 2023; Tsay, Kim, et al., 2022). However, comparing experiment 1, where participants used a movement strategy, and experiment 2, where participants were instructed to aim directly at the movement target, we found that implicit aftereffects were stronger when adaptation was purely implicit, i.e., in experiment 2. This finding is in line with several previous studies that report a negative relationship between implicit adaptation and strategic re-aiming (Albert et al., 2022; Benson et al., 2011), or between implicit adaptation and experimental manipulations which draw attention away from visual feedback (Taylor & Thoroughman, 2007; Tsay, Haith, et al., 2022).

However, implicit adaptation likely depends not solely on changes to proprioception. Patients with cerebellar ataxia exhibit reduced proprioceptive precision compared to healthy controls (Bhanpuri et al., 2013; Weeks et al., 2017a, 2017b), however, they adapt less than controls. Both the reduction in adaptation, and in proprioceptive precision, are attributed to an impairment of internal predictive models in these patients. The perception of limb position, and its relationship to implicit motor adaptation, therefore appear to be more complex, and require further studies.

### Limitations and future directions

Our study was partly motivated by recent reports that highlight a role of proprioception in implicit motor adaptation (Tsay, Kim, et al., 2022). However, our sense of limb position is complex, and reflects various sources of information, including expectation about the intended action, efferent information, proprioception, and vision (Proske & Gandevia, 2009). Here, participants reported hand positions after active movements, blending information from multiple sensory sources. To assess the influence of intermodal attention on proprioception specifically, future studies may include a condition in which hand localization follows passive movements.

We consistently observed a decrease of hand localization precision when participants reported their perceived hand position using a visual pointer. However, when hand position was reported using the 2AFC method, our observations regarding precision were mixed, that is, we found a decrease in localization precision when participants used an aiming strategy to compensate for rotated cursor movement (experiment 1) but not when they ignored the rotation and aimed directly at the target (experiment 2). We hypothesized that participants used the remembered target location as a proxy for hand location during localization. We would expect that such an effect of target location was facilitated in experiment 2, where spatial attention was not divided between two targets, compared to experiment 1 (see also **Supplementary Analysis S4**). Due to its high ecological validity, future studies could use gaze tracking to examine the influence of spatial attention on localization precision. An alternative method to reduce the reliance on the movement target as a proxy for hand localization would be to use targets that are less spatially confined, e.g., arc-shaped targets. We have shown that hand localization precision decreased during adaptation even when hand localization was cued beforehand (experiment 3). The usual point-shaped targets could thus be used for reaching movements during adaptation or washout, and an arc-shaped target could be used for movements immediately before a localization, minimising any influence of the target on localization.

To assess variability of perceived hand position as a proxy of proprioceptive precision, we would ideally examine hand localization variability during a single, extended adaptation period of many trials, followed by an extended washout period, i.e., using a classic design of motor adaptation studies. Computing variability requires repeated measurements. However, during early motor learning, movements and perception change rapidly (Ruttle et al., 2018, 2021). To circumvent this non-stationarity, which could potentially inflate perceptual variability, we decided to probe perception once during early adaptation, and repeat this early adaptation stage many times. However, adaptation may change with repetition and thus affect perceptual measures. For example, the rate of implicit learning changes during the repeated presentation of a visuomotor rotation (Hadjiosif et al., 2023; Yin & Wei, 2020). Additionally, it is conceivable that adaptation and washout cycles in later stages of the experiments were influenced by memory retrieval of errors experienced earlier (Haith et al., 2015; Morehead et al., 2015). Indeed, we found that participants adjusted the instructed re-aiming strategy in adaptation cycles during the later stage of experiment 1, based on the cursor errors they experienced in the beginning of the experiment, which may in turn interact with implicit adaptation (Albert et al., 2022; Miyamoto et al., 2020), and thus influence perception of hand position. In future studies we plan to examine confidence measures of perception (Evans et al., 2022; Olawole-Scott & Yon, 2023) as a potentially faster, more direct alternative to variability of perceptual reports, which may enable assessment of perceptual precision within a single short period of adaptation.

### Conclusion

We report for the first time that the precision of hand localization decreases during early adaptation to a perturbation of visual movement feedback. This reduction is likely caused by focussing attention on visual vs. proprioceptive information during strategy-based motor adaptation.

## Acknowledgements

We thank Denise Henriques for helpful feedback on the study conceptualization and the manuscript. We thank Izel Avci and Melina Trigoussis for support with data acquisition. We are grateful to Richard Ivry for helpful feedback on the manuscript and the members of the Cognition and Action Lab for helpful discussion.

## Author Contribution

M.W., Conceptualization, Data acquisition and curation, Formal analysis, Funding acquisition, Validation, Investigation, Visualization, Methodology, Writing – original draft, review and editing; M.-P.S., Supervision, Conceptualization, Funding acquisition, Validation, Investigation, Methodology, Writing – review and editing

## SUPPLEMENTS

### S1 Reduction of cursor error from early to late experiment is not fully explained by reduced implicit adaptation

An alternative explanation for the reduced cursor error in the later stage of experiment 1 could be that implicit adaptation decreases over the course of the experiment. We found that the size of aftereffects, i.e. the first reach in washout, also changed over the course of the experiment (early: 3.63°±2.63°, late: 0.92°±1.34°, t(29)=5.745, p<.001, d=1.049), possibly reflecting decreased error sensitivity due to the frequently changing perturbation during the experiment (Herzfeld et al., 2014). Alternatively, we cannot exclude the possibility that participants anticipated their own aftereffect during washout cycles, similar to expecting an increase in cursor error during adaptation cycles, and re-aimed in foresight. Importantly, the reduced aftereffect at later stages of the experiment cannot fully explain the decline in overcompensation across the adaptation cycle, as aftereffects were still significantly positive, while change in cursor error was significantly negative. We conclude that participants altered the instructed aiming strategy to minimize cursor errors that resulted from implicit adaptation around the cursor target.

### S2 Statistical analyses of localization bias

#### Experiment 1 – Re-aiming task

For the pointer group in experiment 1 we performed a 2x3-rANOVA with the within-subject factors Cycle Type and Reach-number (i.e. 2^nd^, 3^rd^ or 4^th^ reach in a cycle). We found a significant difference for the factor Cycle Type (F(1,14)=4.63, p=.049, *η*^2^=.23, see **Fig. 3a**) as well as a trend for an interaction between the factors Cycle Type and Reach-number (F(1.25,17.47)=3.44, p=0.07, Greenhouse-Geisser correction). Post-hoc t-tests did not show any significant difference between any of the variables. However, there was a trend for a decrease in perceived hand position bias between 2 and 4 reaches during adaptation cycles (t(14)=3.00, p=0.06, Holm correction). We also confirmed a significant difference in perceived hand position bias between adaptation and washout cycles in the 2AFC group (adaptation 1.69°±3.55°, washout -0.74°±4.38°, t(13)=2.27, p=0.02, d=.61, one sided, see **Fig. 3d**).

#### Experiment 2 – Aim-direct task

For the pointer group in experiment 2, a 2x3-rANOVA with within-subject factors Cycle Type (adaptation or washout) and Reach-number (i.e. 2^nd^, 3^rd^, or 4th reach in a cycle) showed a significant effect of Cycle Type, with a stronger bias during adaptation (7.56°±6.79°) compared to washout (1.65°±4.42°; F(1,13)=41.28, p<.001, *η*^2^=.63). There was no effect of Reach-number(F(2,26)=.20, p=.82), nor an interaction between both factors (F(2,26)=.35, p=.71). In the 2AFC group, hand position reports exhibited a greater angular error during adaptation cycles compared to washout cycles (adaptation: 5.19°±5.16°, washout: 1.74°±3.81, t(13)=4.62, p<0.001, d=1.24, **Fig. 4b**).

#### Experiment 3 – Cued localization

We performed a 2x2-mixed-ANOVA with within-subject factor Cycle Type (adaptation or washout) and between-subjects factor Group (2-target group or 1-target group). We found a significant effect of Cycle Type (adaptation: 9.62°±7.41°, washout: 3.97°±6.91°, F(1,27)= 62.04, p<0.001, *η*^2^=.14, **Fig. 5b**), but no significant effect of Group (F(1,27)=.32, p=0.58), nor an interaction between Cycle Type and Group (F(1,27)=.07, p=0.79).

### S3 Validity of localization variability increase in early visuomotor adaptation

We conducted several control analyses to exclude the possibility that the observed increase in variability of hand localization was an artefact of any systematic changes in hand localization across the experiment, or in movement endpoints between adaptation and washout.

First, we compared bias during the first 30 adaptation cycles and the last 30 adaptation cycles (pointer group). We found no effect of experiment stage on bias (t(14)=1.33, p=.207). However, a Bayesian paired t-test did not yield any convincing evidence for either hypothesis (BF_10_=.547). We thus corrected for any drift in angular localization errors over the experiment by detrending hand localization reports using linear regression (separately for each cycle type and target, see **Fig. 6a**). The IQR of corrected perceptual reports remained significantly different for adaptation and washout cycles even after detrending (adaptation: 9.81°±4.13°, washout: 5.78°±1.75°, t(14)=4.77, p<.001, d=1.23).

We also tested directly whether the variability of hand localization during adaptation cycles differed between early and late stages of the experiment (first 30 vs. last 30 cycles). We did not find any difference between the early and late stage of the experiment (early: 9.31°±4.07°, late: 9.08°±3.43°, t(14)=.33, p=.75). A Bayesian paired t-test provided moderate evidence for the null hypothesis (BF_10_=.28). There was a decrease of hand localization variability from the early to the late stage of the experiment during washout cycles (early: 6.33°±2.22°, late: 5.08°±2.01°, t(14)=2.63, p=0.02, d=.68).

Finally, it is possible that participants based their estimate of hand location (partly) on the remembered location of the target. If the difference between the actual hand position and the remembered visual target exhibited higher variability during adaptation cycles, compared to washout cycles, any such bias in hand localization towards the target could have inflated variability in hand perception in adaptation cycles. Indeed, we found evidence that hand positions became systematically more variable during adaptation (11.32°±5.67°), compared to washout (3.80°±.64°, t(28)=7.08, p<.001, d=1.32). However, we could rule out the possibility that the observed increase in hand localization variability during adaptation merely reflected an increase in hand position variability. Specifically, we matched the hand position variability during localizations between adaptation and washout cycles in the pointer group. This was achieved by iteratively removing trials with extreme hand positions in adaptation, and average hand positions in washout (85 trials removed on average across subjects (median; range 48-127), 66 trials remaining (median; range 32-84)). We then recomputed the variability of hand localization for the remaining trials, and performed a 2x2-rANOVA with the within-subject factors Cycle Type (adaptation or washout) and Type of variability (hand position or localization). We found a main effect of Cycle Type (F(1,14)=11.3, p=.005, *η*^2^=.13), and, importantly, an interaction between Cycle Type and

Type of variability (F(1,14)=12.7, p=.003, *η*^2^=.16, see **Fig. 6b**). Post-hoc t-tests showed that there was no significant difference between hand position variability between adaptation and washout cycles (adaptation: 6.51°±1.56°, washout: 6.63°±1.65°, t(14)=.2, p=.88, Holm correction; BF_0-_=11.40, one-sided), however, hand localization variability was still higher in adaptation, compared to washout (adaptation: 8.61°±3.92°, washout: 5.90°±2.14°, t(14)=2.97, p<.001, Holm correction).

**Fig. 6:**
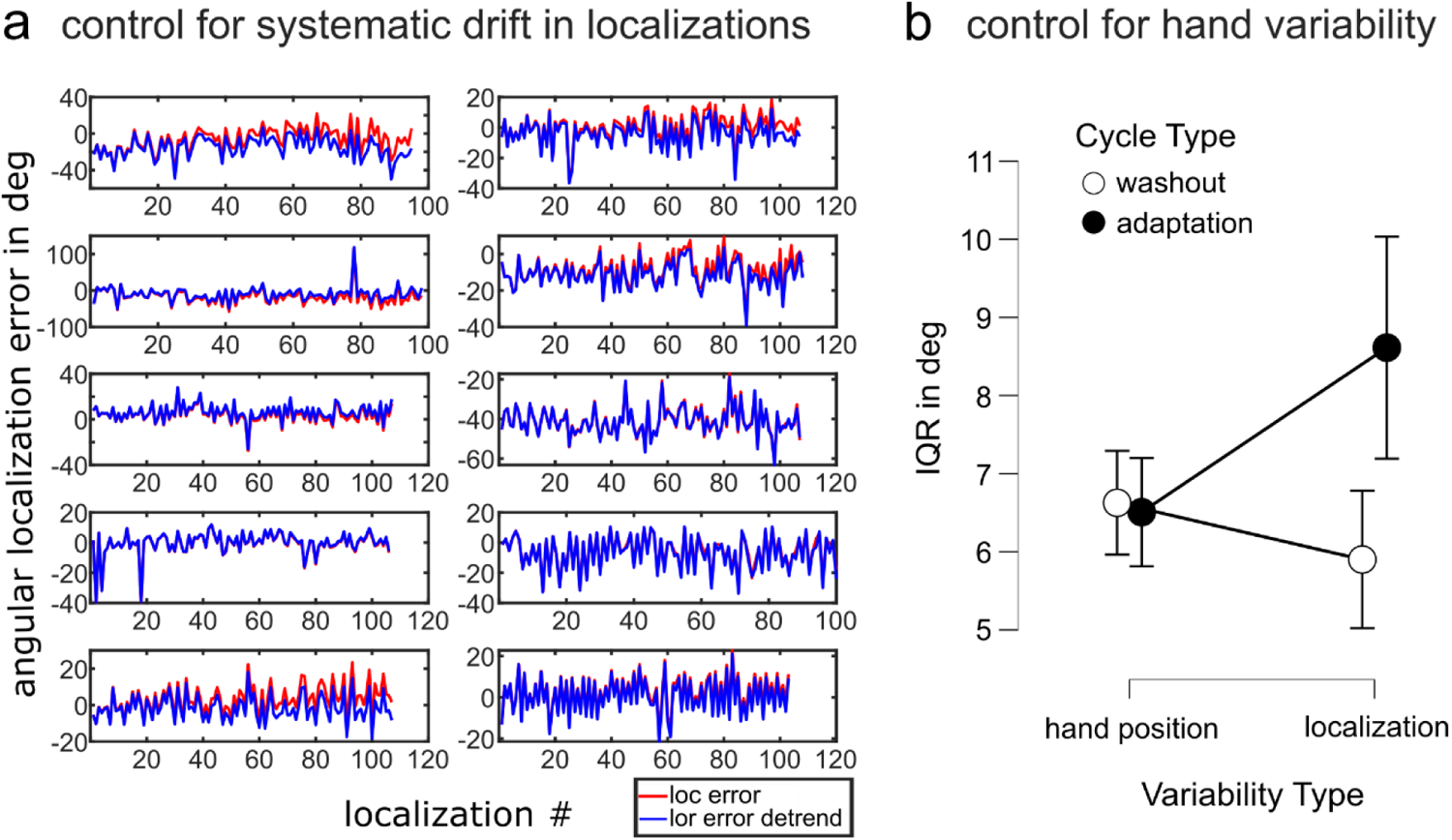
Control analyses (a) Potential systematic drift in localization errors (here for adaptation cycles, red curves) was removed by detrending using linear regression (corrected curves in blue, different panels show different exemplary participants). Few participants showed drift (e.g. top left or bottom left). However, the observed increase in localization variability persisted after removing any linear change in hand localization error across the experiment (b) When hand position variability during localizations (left) was matched between adaptation and washout (black and white dots, bars indicate standard error of the mean), localization variability (right) in adaptation cycles was still higher, compared to washout cycles (N=15).

### S4 Discrepant hand localization variability between pointer and 2AFC groups may be attributed to enhanced emphasize of remembered target location

To explain the discrepant findings for hand localization variability between the pointer group and the 2AFC group, we considered a potential role of remembered target location. In the pointer group, the presentation of the pointer for localization, close to the home position, likely resulted in a large shift of gaze away from the hand target location, towards the pointer. Similarly, both groups of experiment 1 likely focused on the task-relevant cursor target at the beginning of a localization trial, i.e., away from the hand target. However, subjects in the 2AFC group of experiment 2 likely focused gaze on the location of the hand target from the beginning of a localization trial on (given that there was no other task-relevant visual stimulus) which likely enabled them to remember the target location more precisely. The remembered location of the hand target may thus have influenced hand localization reports more strongly in the 2AFC group of experiment 2 than in the pointer group, and in both groups of experiment 1, whose gaze away from the hand target location may have instead emphasized proprioceptive information for localization.

To test this idea, we compared two generalized linear models predicting the reported hand position during adaptation cycles. The full model explained position reports based on two predictors, i.e., the probe relative to the hand and relative to the target. The sparse model, on the other hand, explained position reports based on the probe relative to the target only. Model comparison using Akaike information criterion (AIC) showed that the sparse model explained hand reports better than the full model in 5 out of 14 participants of the 2AFC group in experiment 2, reducing AIC by >1.5 (indicating that the sparse model is more than twice as likely as the full model). However, when performing the same analysis with data from the 2AFC group in experiment 1, where participants likely attended the (task-relevant) cursor target at the beginning of localization trials, de-emphasizing the hand target location, the sparse model described individual data better than the full model in none of 14 participants.

Furthermore, we compared the time between the onset of the probe stimulus and the submission of the psychophysical report during adaptation and washout cycles between 2AFC groups in experiment 1 and 2. The rationale was that a shift of gaze costs time. As the 2AFC group in experiment 1 could use their gaze position as a proxy for hand localization during washout (during which they likely focussed on the hand target), but not during adaptation (during which they likely focussed on the cursor target), we expected that localization reports took more time during adaptation. On the other hand, the 2AFC group in experiment 2 could use the same localization strategy in both cycle types, as no (large) shift of gaze was needed in both cycle types. In the 2AFC group of experiment 2, we therefore expected to find no difference between localization durations during adaption and washout cycles. We performed a 2x2-mixed ANOVA with within-subject factor Cycle Type and between-subject factor Experiment, and found a significant interaction between both factors (F(1,26)=4.51, p=.04, *η*^2^=.15). There was a trend for a difference between Cycle Types (F(1,26)=4.15, p=.05, *η*^2^=.004). Post-hoc t-tests revealed a significant difference between localization duration in adaptation and washout cycles in experiment 1 (adaptation: 1912±904 ms, washout: 1710±829 ms, t(13)=2.94, p=.04, Holm correction), which may reflect additional time costs due to a large shift of gaze during adaptation. On the other hand, a Bayesian t-test provided evidence in favour of the null hypothesis that localization durations in the two cycle types in the 2AFC group of experiment 2 were equal (adaptation: 1844±696 ms, washout: 1848±785 ms, BF_10_=0.271). While hand localization reports were not speeded, and response time difference therefore have to be interpreted with caution, this nevertheless further supports the idea that the different localization methods may account for discrepant findings observed for hand localization precision in the two groups of experiment 2.

### S5 Post-hoc comparison of hand localization variability across experiments

To draw conclusion about the contributions of different motor learning mechanisms to the observed perceptual changes, we compared the increase in hand localization variability during adaptation, relative to washout, across the three experiments. For this, we conducted a one-way ANOVA that included all participants (i.e., also collapsed across groups with different hand localization methods), and the between-subject factor Experiment. We found that the increase in variability was largest in experiment 1 (exp. 1: 4.06°±3.79°, exp. 2: .69°±2.09, exp. 3: 1.89°±2.38°, F(2,84)=10.41, p<.001, *η*^2^=.20, post hoc tests: exp. 1 vs. exp. 2: t=4.50, p<.001, exp. 1 vs. exp. 3: t=2.90, p=.013). This was also the case when we only included the pointer group for experiment 2, where we observed a significant increase of hand localization variability during adaptation cycles (exp. 2 pointer only: 1.33°±2.03°, F(2,70)=5.70, p=.005, *η*^2^=.14, post hoc tests: exp. 1 vs. exp. 2: t=2.90, p=.014, exp. 1 vs. exp. 3: t=2.78, p=.019).

## Footnotes

1 We excluded one subject from all analyses of hand localization data who did not follow instructions during the hand localization.

2 We excluded one subject of the pointer group from the analysis due to implausibly large angular localization errors during washout cycles. In the 2AFC group, the PSE of two participants could not be computed correctly during adaptation cycles as relative frequency of answering “left” for all probe values fell short of .5. Furthermore, the psychometric curves obtained for washout cycles of two subjects exhibited a very sharp slope, resulting in a JND that was implausible (<0.3°). Thus, we excluded these subjects from the analysis.

